# The SAFE Labs Handbook: Community-Driven Commitments for Group Leaders to Improve Lab Culture

**DOI:** 10.1101/2025.05.27.655799

**Authors:** E Donà, JM Gahan, P Lau, J Jeschke, T Ott, K Reinhard, C Sinigaglia, JL Treur, T Vogl, S Bugeon, L Mariotti, LF Rossi, P Coen

## Abstract

Creating positive and equitable lab environments has become a growing priority for the scientific community and funders of scientific research. Research institutions typically respond to this need by providing mandatory or optional training opportunities for their staff. However, there is no established resource for group leaders to improve their lab culture with concrete action points for implementation. Here, we introduce the SAFE Labs Handbook: a collection of thirty “commitments” which can be verifiably actioned without requiring institutional support. These commitments were collaboratively developed by thirteen group leaders in life sciences, from institutions across eight countries, instigated through the 2024 SAFE Labs workshop. The importance of each commitment has been scored by more than 200 researchers, at various career stages, from more than twenty countries. Even though all commitments were rated as significantly important by scientists from all career stages, implementation rates were notably low (< 25%). Lab members reported higher importance scores than group leaders, with large divergences indicating where group leaders may underestimate the potential impact on lab culture. Indeed, the overall implementation rate was correlated with importance score for group leaders, but not lab members. Strikingly, more than 95% of group leaders said they would consider implementing the handbook commitments. Given the high importance-scores and low-implementation rates, the SAFE Labs Handbook represents a unique, community-driven tool with significant potential to improve lab culture on a global scale.

## Introduction

The importance of creating a positive, fair and transparent working environment in academia has become increasingly evident. Academic environments are associated with impaired mental health, poor work-life balance, and disparities in pay and career progression (Acton et al., 2019; Evans et al., 2018; Guthrie et al., 2018; Kim et al., 2024; Kinman and Jones, 2008; Marck et al., 2024; Morgan et al., 2021; Susi et al., 2019; Wellcome Trust, 2020). Solving these problems requires intervention at all levels of the academic hierarchy, but the scope and feasibility of institutional and educational changes can vary dramatically across countries. Top-down changes at the national or institutional level are slow to enact. Conversely, bottom-up changes, implemented by individuals, can be adopted more rapidly and without administrative delays.

The “research group” is the foundational unit throughout the academic world, whose policies can determine the wellbeing of researchers (Halat et al., 2023). Group leaders are typically responsible for providing training and supervision for their lab members and have agency to determine lab policy and to influence the working environment. There are informative resources with advice for starting a research group (Aly, 2018; Goldstein and Avasthi, 2021; Schmidt, 2006; Somerville et al., 2019), but these typically lack specific action-items to help group leaders create fair and equitable teams. Further, existing resources have not assessed the level of support for their recommendations throughout the academic community. Therefore, an effective, community-driven tool for group leaders to improve lab culture has the potential to have a major impact throughout academia both for mental well-being and academic productivity (Lohela-Karlsson et al., 2018). Such a tool would not eliminate overtly negative behaviour by abusive individuals (Moss and Mahmoudi, 2021), but would instead support group leaders that are committed to improve the culture in their research environment.

No group leader has a universal formula for success, but transparency empowers prospective members to find the environment that matches their needs. A single set of guidelines cannot capture the diversity of personalities, preferences, and research fields represented across group leaders and their lab members. To account for this, existing resources for improving lab culture have focused on clear and transparent communication with lab members (Aly, 2018; Andreev et al., 2022; Martin and Stanfill, 2023; Tendler et al., 2023). These are valuable and insightful materials, but they are often limited to case studies that represent the views of an individual or small group and lack evidence from the community to support their recommendations. Conversely, larger studies may have hundreds of participants, but tend to highlight broader systemic problems in academia, proposing large-scale reforms that cannot be implemented at the level of an individual research group (Acton et al., 2019; Marck et al., 2024; Strategy Group, 2019; Wellcome Trust, 2020).

At the 2024 <u>S</u>tarting <u>A</u>ware, <u>F</u>air, and <u>E</u>quitable <u>L</u>abs (SAFE Labs) workshop, new group leaders from across Europe explored how to foster more positive and equitable lab environments. We identified the need for a set of actionable “commitments” that could be verifiably implemented by individual group leaders without institutional support. Although institutions have an important role in improving lab environments, they can be limited by national policies and lengthy administrative processes. Here, we present the resulting *SAFE Labs Handbook* (Figure 1), a unique tool to empower group leaders to create more positive and equitable research environments. The thirty commitments in this document have been evaluated and scored by more than 200 researchers, including group leaders, postdoctoral researchers, PhD students, and staff. The complete handbook, along with explanations of each commitment and example text, can be found here: https://safelabs.info/home/safe-labs-handbook/.

**Figure 1.**
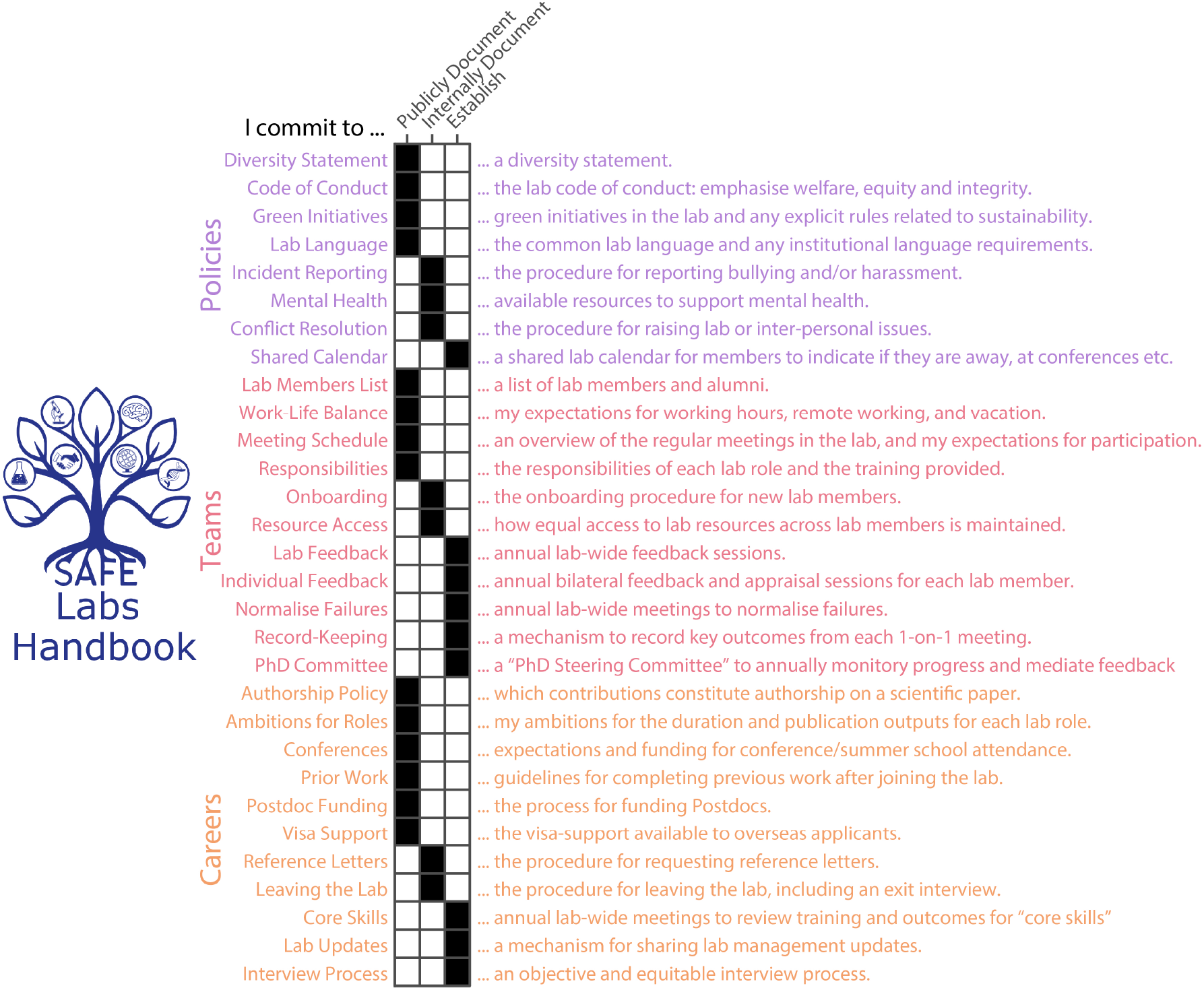
SAFE Labs Handbook Commitments. The SAFE Labs Handbook logo (left) and summary of its commitments (right), arranged by category. Matrix indicates (black) whether each commitment should be publicly documented, internally documented, or established. The complete handbook, along with explanations of each commitment and example text, can be found here: https://safelabs.info/home/safe-labs-handbook/.

## Results

### 2024 SAFE Labs Workshop

The 2024 SAFE Labs workshop provided a unique opportunity to discuss how to create aware, fair, and equitable research environments. The workshop was attended by thirteen new group leaders (four organizers and nine attendees) from eight European countries, working in different disciplines across the biosciences (Supplemental Figure 1c,d). Although cultural problems in academia are wide-ranging, the workshop focused on how group leaders could improve their lab environment without institutional support, with discussion topics and formats chosen using applicant data (Supplemental Figure 1a,b). The feedback on this workshop was overwhelmingly positive, with 100% of attendees stating that it exceeded expectations and agreeing that *SAFE Labs was a unique meeting that fulfilled an important niche* and that they would *change practice in their labs because of attending SAFE Labs* (Supplemental Figure 1e-g).

The SAFE Labs workshop highlighted two key barriers to improving lab culture as a new group leader. First, existing resources provide general advice rather than actionable steps that can be verifiably implemented (Aly, 2018; Andreev et al., 2022; Strategy Group, 2019; Tendler et al., 2023). Second, academia does not normalise the documentation of lab policy, with group leaders often relying on *word-of-mouth* to educate new lab members. When documentation does exist, key information is often limited to existing lab members, and not prospective applicants. SAFE Labs attendees agreed that the “best” lab environment was dependent on the individual. However, labs could universally benefit from documenting key policies to minimise expectation mismatch between lab members (existing and prospective) and the group leader. To achieve this goal, attendees proposed to create a “SAFE Labs Handbook” with 100% of attendees agreeing (through an anonymous survey) to contribute. The handbook was designed to both provide actionable steps for group leaders to improve their lab environments, and to normalize the process of documenting key lab policies.

### SAFE Labs Handbook

The SAFE Labs Handbook comprises thirty commitments spanning three broad categories: SAFE Teams, Policies, and Careers (Figure 1). Each “commitment” is an actionable statement that asks a group leader to *publicly document, internally document*, or *establish* a policy in their laboratory (Figure 1b). *Publicly* documented information should be visible to anyone (e.g. posted on the lab website). *Internally* documented information need only be visible to existing lab members (e.g. posted in a password-protected lab manual or wiki): this may be information that is sensitive, or only relevant to individuals that are current members of the lab. *Establish* indicates a commitment that requires a change at the level of lab management, such as instantiating a new annual meeting. Commitments were generated through a distillation of discussion notes at the 2024 SAFE Labs workshop by the thirteen participants (see Methods). *Publicly* vs *internally* documented commitments were established through anonymous vote, with the decision deferred to the community-survey if there was no consensus (10% < score <50% in favour, Supplementary Figure 1h).

Commitments were chosen to be actionable and verifiable to facilitate implementation and accountability. Each commitment derived from collated documentation from the SAFE Labs Workshop. We identified prominent issues that warranted inclusion in the handbook. For example: expectations around working hours and vacation are unclear to prospective and current lab members. We then agreed on the form of the commitment that made it *actionable, verifiable* and *minimally prescriptive*. “I commit to supporting a healthy work-life balance” would *not* qualify because the commitment cannot be verifiably implemented. Instead, the analogous entry in the handbook states that a group leader commits to…

> *“… publicly document* expectations for working hours, remote working, and vacation.*”*

This is actionable (requires specific action from the group leader), verifiable (implementation can be evidenced), and does not mandate any particular policy—it instead requires that the policy is documented so that expectations are clear to all current and prospective lab members. We therefore provide only *details, suggestions*, and where appropriate, a freely editable *template* statement for each commitment. We also encourage the growing community to add examples of their statements, to help new labs develop effective policies tailored to their environment. As an illustration, here is the current entry for the above commitment:

#### Details

> *Lab rules for working hours should be clear to avoid conflicts or misunderstandings, and clear expectations for working hours can increase equity between lab members. Group leaders need to ensure that lab members feel safe to balance their work in the lab with their life outside it. Policies regarding work hours, remote working, and vacation should be explicitly included*.

#### Suggestions

– *Should notice of holidays be given and how?*
– *Are there core-working hours (typically less than the full working hours)?*
– *Should lab members schedule messages if sent outside of working hours?*
– *What times are appropriate times for scheduling meetings?*

#### Template

> *I am committed to creating a healthy work environment for all lab members. I anticipate all lab members taking a minimum of [institution’s] prescribed days of annual leave. “Minimum” because if experiments/conferences necessitate working on a weekend, I support lab members taking time off to compensate. I hope to schedule all meetings within [institution’s] “core” work hours, and will refrain from sending, or answering, non-urgent emails/messages outside of work hours*.

> *Full-time lab members should work “onsite” at least four days a week. I believe regular onsite presence is important to maintain the lab community. However, I will support intermittent periods of remote work when, for example, traveling/visiting family abroad or writing up a thesis/grant*.
>
> To assess the need for each handbook commitment, we surveyed over 200 academics from more than 20 countries (4 continents, Figure 2). Survey participants rated the importance of every commitment and stated whether it was already implemented in their environment. For a subset of commitments, participants were also asked whether information should be *internally* or *publicly* documented. The most common respondent career stage was “group leader” (51%, n = 105), suggesting a strong drive to improve lab culture at the senior level, with the remaining 49% split amongst postdocs (25%, n = 57), PhD students (21%, n = 46), and research support staff (7%, n = 16) (Figure 2a). Further, more than 30% of respondents (over 40% of non-PhD student respondents) had been in their role for over 4 years—demonstrating that both new and experienced members of the community engaged with our survey (Figure 2b). Participants represented institutions in 24 countries and worked in a range of fields (Figure 2c,d). Although these demographics are influenced by organiser bias, no single country represented more than 25% of participants. Overall, participant demographics indicate broad engagement with the SAFE Labs Handbook.

**Figure 2.**
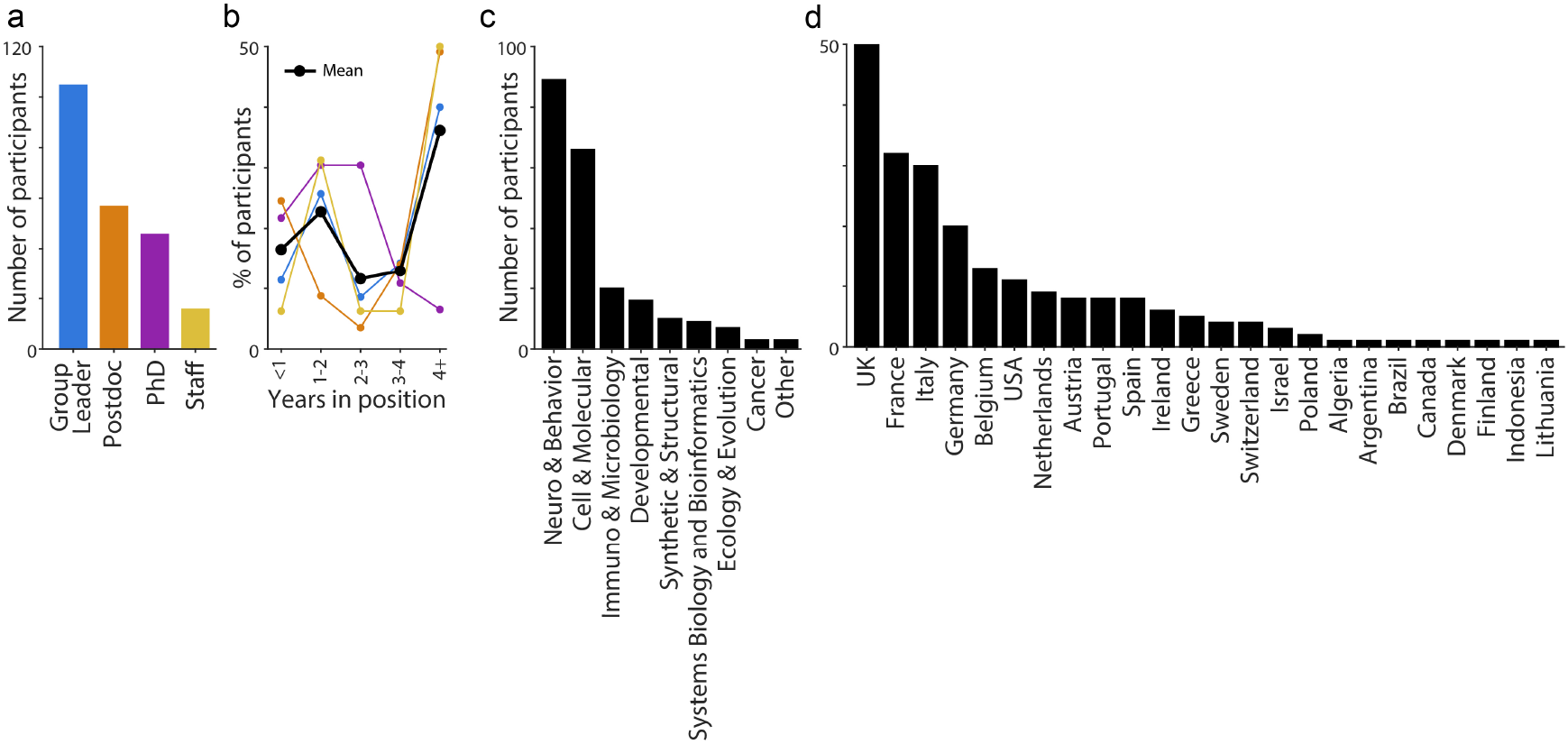
Demographics of Handbook Survey Respondents. (**a**) Number of respondents to the SAFE Labs Handbook survey, sorted by employment role. (**b**) Fraction of respondents as a function of seniority in each role from (a, color). Black line indicates the average across all respondents. (**c**) Number of respondents sorted by research field. (**d**) As in (c), but by country of employment. For all panels, n = 105 Group Leaders, 57 Postdocs, 46 PhD students, and 16 Staff.

### Community Survey

All SAFE Labs Handbook commitments were deemed important by participants at all career stages. When ranking commitments from 1 (useless) to 5 (critical), participants scored all commitments above 3, even when answers were subdivided by career stage (Figure 3a-b) or country (Supplementary Figure 3a-d). Strikingly, despite this universal agreement on importance, commitments were implemented at a rate of only 25% across labs (Figure 3c-d), illustrating the need for a document like the SAFE Labs Handbook to bridge this gap. The mean importance score amongst respondents was not significantly impacted by their years of experience (Supplementary Figure 2). Across the 5 countries with the most respondents, relative importance of commitments and implementation-rates were broadly consistent, with the UK and Italy reporting the highest and lowest implementation rates (Supplementary Figure 3a-d). There was a strong correlation (R = 0.67, p < 0.001) between implementation and importance of a commitment, as rated by group leaders (Figure 3e). This suggests that group leaders prioritize commitments they perceive as more important, but this may also reflect a confirmation bias. The same correlation was also significant when separated by country (Supplementary Figure 3e-I).

**Figure 3.**
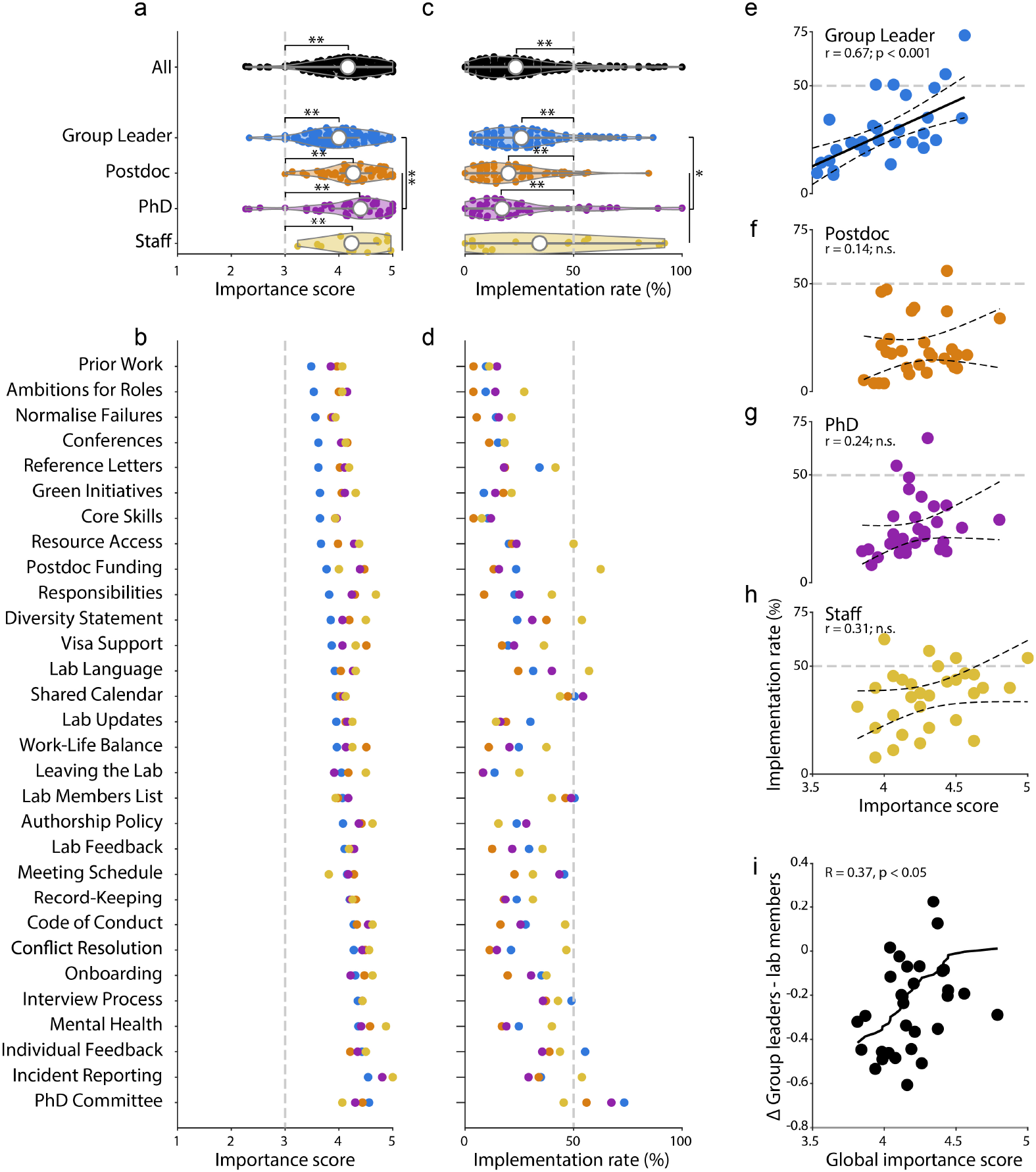
Differential commitment rating between lab members and group leaders. (**a**) Distribution of mean importance score across commitments, for all survey respondents (black), and for each career stage: Group Leader (blue), Postdoc (orange), PhD student (purple), and Staff (yellow). Violin plot shows median (white dot) and range. All career stages rated the handbook commitments as significantly important (i.e. above the gray dotted line, ∗∗ = p < 0.001). The importance scores from group leaders were significantly lower than lab members (Mann–Whitney U test, ∗∗ = p < 0.001). (**b**) Mean importance score for each commitment and career stage, sorted in ascending order based on Group leaders’ answers. (**c**) As in (a), but for the implementation rate. All career stages reported low levels of implementation, significantly below 50% in most cases (gray dotted line, ∗∗ = p < 0.001). The implementation rate indicated by group leaders was significantly higher than lab members (Mann–Whitney U test, ∗ = p < 0.05). (**d**) As in (b), but for the implementation rate. (**e**) Relationship between mean importance and mean implementation rate for each commitment as scored by Group Leaders. The significant correlation was captured by a robust linear fit (black, dotted confidence intervals, p < 0.001) (**f**) As in (e), but for postdocs (p > 0.05). (**g**) As in (e), but for PhD Students (p > 0.05). (**h**) As in (e), but for Staff (p > 0.05). (**i**) The difference in importance score between group leaders and lab members is significantly correlated with the global importance across all groups (black, Spearman rank R = 0.37, p < 0.05).

Lab members rated commitments as more important, and less established, than group leaders. Commitments had a higher importance rating when scored by lab members than group leaders (Figure 3a-b, p < 0.01, MWU test), suggesting that group leaders systematically underestimate the importance of these commitments to their lab members. Surprisingly, lab members also reported a lower rate of implementation than group leaders (Figure 3c-d, p < 0.05, MWU test). This may be because group leaders overestimate the completeness of their implementation, or that lab members are unaware of the policies—reinforcing the importance of communicating lab policy through a written medium. Unlike for group leaders (p < 0.001, robust linear fit), we did not observe a significant correlation between commitment importance and implementation for lab members (Figure 3f-h), suggesting a disconnect between the actions taken by group leaders, and those that would be most valuable from the perspective of their group. Supporting this, we found that increased divergence between group leaders and lab members was significantly correlated with a higher global importance score (Figure 3i, p < 0.05, Spearman correlation).

Lab members and group leaders supported internal, rather than public, documentation of lab policies. When SAFE Labs Handbook authors did not agree on whether a commitment should be *publicly* or *internally* documented (Supplementary Figure 1h), survey respondents were asked for their input. Group leaders indicated a broad preference against *public* documentation (*public* preference ranged from 9-26%), while lab members were more supportive (12-75% *public* preference) (Supplementary Figure 4). These results suggest that while *public* documentation is considered valuable to some lab members, it is most important the information *is documented*. This informed our plans for the handbook (see Discussion).

**Figure 4.**
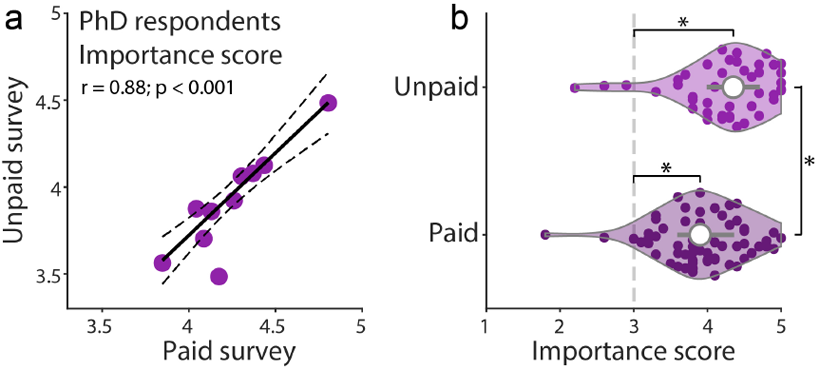
Highly correlated survey outcome between paid and voluntary survey respondents. (**a**) Relationship between mean importance rating of 10 commitments (subset chosen for the paid survey) by PhD students from the paid and unpaid survey. The significant correlation was captured by a robust linear fit (black, dotted confidence intervals, p < 0.001). (**b**) Distribution of importance rating averaged across commitments for respondents of the paid and unpaid surveys (Mann– Whitney U test, p < 0.01). n = 46 voluntary and 64 paid PhD respondents.

Responses of voluntary participants were correlated to those of paid participants. Our survey results could be biased by sampling individuals willing to volunteer their time to complete the survey. To test the general applicability of our results, we therefore ran a paid survey, asking current PhD students to rate a subset of 10 commitments (see Methods). In support of our initial survey, there was a strong correlation (p < 0.001, robust linear fit) with unpaid participants when considering the relative importance across commitments (Figure 4a), although importance scores were marginally lower for this paid cohort (Figure 4b).

Respondents supported the implementation of the SAFE Labs Handbook in their group. More than 50% of group leaders, and 70% of lab members (both unpaid and paid participants), supported full implementation of the handbook in their lab environment (Figure 5, Supplementary Figure 5). Critically, less than 5% of respondents were unsupportive of the handbook. Instead, a significant fraction of respondents indicated “maybe”—often citing institutional limitations or preferences for internal documentation (see Discussion). Given the verified importance of these commitments, this level of implementation has the potential to generate systemic change throughout the academic community.

**Figure 5.**
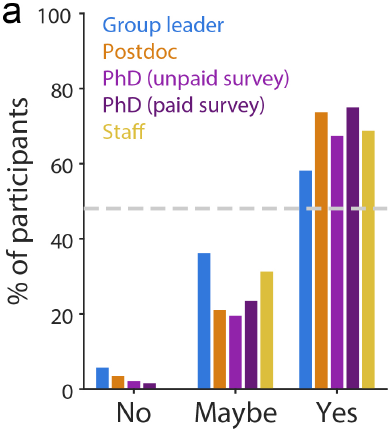
High support for handbook implementation across all respondent groups. Fraction of respondents from each survey (paid and unpaid) in favor of implementing the handbook in their environment. For all panels, n = 105 Group Leaders, 57 Postdocs, 46 PhD students, 64 paid PhD students and 16 Staff.

## Discussion

To provide a tool for group leaders to create more positive and equitable lab environments we developed the SAFE Labs Handbook, a collection of 30 commitments which can be implemented by individual labs without institutional support. Co-authored by 13 early-career group leaders from across Europe, these commitments were subsequently assessed by more than 200 participants from more than 20 countries. All commitments were considered important by both group leaders and lab members, and 95% of group leaders (50% *yes*, 45% *maybe*) indicated their willingness to implement the handbook in their lab. The initiative also attracted 132 of subscriptions to the SAFE Labs mailing list to engage with future workshops and initiatives.

A cornerstone of the SAFE Labs Handbook is the recognition that each group leader has a distinct approach to management. Thus, *commitments do not specify how a lab should be managed*, and different approaches are often equally valid, and appealing to different lab members. We do not envision that this tool will resolve all issues, or stop overt abuses of power (Moss and Mahmoudi, 2021). The handbook instead focuses on improving transparency and written documentation of lab policy to minimize mismatched expectations between the group leader and their current or prospective lab members. Existing materials have also emphasised this approach (Aly, 2018; Andreev et al., 2022; Somerville et al., 2019; Tendler et al., 2023), but have not distilled their recommendations into verifiable commitments, and have not quantified the level of support for those recommendations throughout the academic community. The importance of lab culture, and the motivation behind the selection of these commitments formed a major component of the SAFE Labs workshop in 2024. We have chosen to omit this broader discussion here to maintain focus on providing a practical tool for group leaders, especially since the survey results demonstrate that the importance of these issues is already evident throughout the community.

The SAFE Labs Handbook commitments are widely recognized as important but are implemented in a minority of labs. Despite all participant groups agreeing on the *importance* of the commitments, the rate of *implementation* was notably low (< 25%). We believe this percentage may even be an overestimate, as respondents included lab members from the authors of this document, where implementation is more probable. This gap between perceived importance and actual implementation demonstrates the value that the SAFE Labs Handbook can bring to the community. Group leaders are already convinced of the importance of these commitments, the handbook must simply work to increase *awareness*, accompanied by example documentation to ease implementation. Although implementation rate and importance were strongly correlated for group leaders, this correlation was not significant for lab members, suggesting that group leaders could better align their efforts with those of their lab members.

Survey respondents reflect significant sampling biases, but correlate with the opinions of an independent cohort of paid participants. The predominance of survey respondents from the UK, France, and Italy, as well as the fields of Neuroscience and Cellular biology, reflect the composition of the authors of this manuscript, as well as biases in grant mechanisms and the willingness of participants to volunteer their time (see Methods). Our results were strongly correlated with data from a separate pool of paid participants, indicating that these results reflect opinions of the wider academic community. Nevertheless, we acknowledge that the handbook would benefit from an increased diversity of voices. Indeed, with more data, differences in implementation-rate or perceived importance across countries could serve as important indicators that changes need to be made at the institutional or national level to support group leaders.

The SAFE Labs Handbook is a dynamic resource document that will evolve through community-driven feedback and contribution. Because the SAFE Labs Handbook is innovative, it remains untested. Our survey data establishes the need for this tool, and the importance of the existing commitments. However, there remains significant scope for both revision and expansion, both from feedback as labs implement the handbook, and from further dedicated workshops. The handbook repository already includes a growing collection of real-world examples of statements from labs implementing the handbook, which will help new labs develop their effective policies. Moreover, future workshops, like the 2025 SAFE Labs workshop, will focus in-part on proposing changes to the handbook via the lens of a new cohort of group leaders and organisers (https://safelabs.info/).

In addition to quantifiable metrics, our survey presented participants with the opportunity to comment on individual commitments, and on the complete handbook. We address the four most common concerns below:

### Concern 1: A commitment is covered at the institutional level

This was most common with respect to aspects of employment contracts (e.g. working hours, holiday). We now clarify that “documenting” could involve linking an institutional policy and stating that it reflects your expectations for lab members. However, exceptions to institutional policies should be clarified (e.g. remote-working, experiments/conferences on weekends). Group leaders can provide an anonymous feedback channel for lab members to highlight any edge-cases that remain unclear.

### Concern 2: The situation is too individualistic, so documentation would be meaningless

This was most common when commitments impacted individual projects (e.g. authorship, resource management, funding of positions). We argue that this perspective (that the situation is too individualistic) also reflects an expectation that should be documented and would still be informative for lab members. We reiterate that the handbook is *not* prescriptive, and that a commitment would be satisfied by stating that no general lab policy exists.

### Concern 3: I do not feel comfortable publicly documenting this information

Several group leaders indicated this was the primary reason that they would “maybe” implement the handbook. As group leaders, we understand this perspective, particularly given that publicly documented material in academia is atypical. We maintain that public documentation is a valuable resource for prospective lab members, and this in turn increases the likelihood of group leaders recruiting suitable candidates for positions. This is evidenced by the higher enthusiasm for public documentation amongst lab members. However, to encourage broader adoption of the handbook, we have removed the *requirement* for public documentation from many commitments.

### Concern 4: Implementing these commitments is too much work

This criticism referenced two features: meetings and documentation. Although the handbook encourages group leaders to document regular meetings and expectations, complete implementation of the handbook only requires *three annual meetings* (normalising failure, lab-wide feedback, and core-skills development). This would minimally impact the annual calendar of most labs. Conversely, the documentation required represents a significant workload, exceeding the norm in academia. However, we envision this time will be recouped by pre-empting questions from lab members and reducing conflicts that arise from expectation mismatch or inter-personal differences. We provide concrete templates and suggestions for each commitment to simplify implementation. Furthermore, we have a growing list of community examples from group leaders that have implemented the handbook (already a minimum of two for each commitment). We anticipate implementation times to continue shrinking as this repository expands, and group leaders can select an example that mirrors their own policies.

We envision the SAFE Labs Handbook as a critical tool for group leaders, as well as an opportunity to advertise their commitment to creating more positive and equitable academic environments. We took inspiration from the Laboratory Efficiency Assessment Framework (LEAF), a pioneer in the related space of lab sustainability. Laboratories are given awards based on their sustainability practice (bronze, silver, or gold) and engaging with this process is now a requirement for certain funding sources. We propose to develop an analogous system for the SAFE Labs Handbook, where group leaders indicate the commitments they have implemented and receive an award (weighted by importance, and *public* vs *internal* documentation). This award can be featured on the lab website, referred to in funding applications, and assessed at an institutional level as a quantifiable metric to track improvement (or lack thereof) across years. Compliance with The SAFE Labs Handbook will therefore serve to advertise the importance of lab culture to prospective scientists, at both lab and institutional levels.

## Acknowledgment

The SAFE Lab initiative was funded by the UCL Global Engagement Fund (to SB, LM, LFR and PC). The workshop was hosted under the patronage of Scuola Normale Superiore and the sponsorship of Professor Franco Ligabue. We would also like to thank the 224 participants who volunteered their time to complete the survey. We thank Sara Saab (Prolific) for facilitating the paid survey and Prolific for providing credit to partially cover these costs.

## Contributions

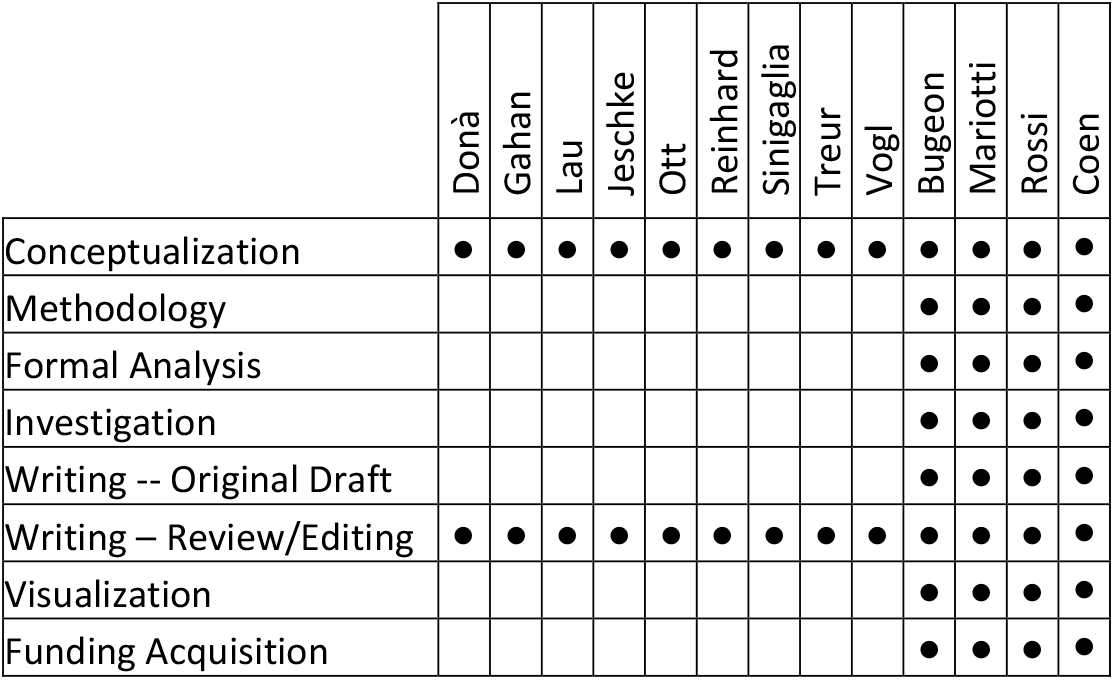

## Methods

### SAFE Labs 2024 Workshop application process

The workshop was advertised through a combination of social media posts and targeted emails to newly appointed group leaders across Europe, within 3 years of starting their lab through awarded grants (e.g. the ERC starting grant). The application included 14 questions, spanning demographics, motivational statements, and potential discussion topics (Supplementary Table 1). The data and ideas from the application were analysed to shape the SAFE Labs workshop (Supplementary Figure 1a-b). 9 participants were selected from a total of 21 applicants based on blind-scoring by the organisers, weighted by the need to maximise diversity across countries and scientific fields.

### SAFE Labs 2024 Workshop Format and Handbook Preparation

The workshop comprised 6 sessions, based on topics highlighted by applicants. Each session had three phases: an introductory presentation, small group discussions, and a joint roundtable. The introductory presentation was prepared in advance and delivered by one of the applicants. For group discussions, participants were randomly divided into three subgroups, each with a randomly assigned group leader and note-taker, to discuss three questions on the current topic: What are the biggest issues facing new labs? What can new PIs do to tackle/prevent these issues? What could institutions do to support new PIs? During the joint roundtable, subgroups convened to summarise and discuss the main conclusions from each subgroup.

### Workshop Exit Survey

After the workshop, the organizers solicited feedback from attendees through an anonymous exit survey, consisting of 22 questions. Participants were asked to indicate their level of agreement with selected statements, to score the session topics and formats, and the logistical aspects of the meeting (Supplementary Table 2, Supplementary Figure 1e-h).

### SAFE Labs Handbook Voluntary Survey

The voluntary survey was designed to ascertain the level of support for the SAFE Labs Handbook from the research community. The survey was advertised through social media, by soliciting scientific organisations and publications, and direct emails to grant awardees and institutional contacts. In particular, the survey was emailed to recipients of the ERC starting grant from the year 2021 to 2024, and other national early career grants (e.g. Italian PRIN under 40, Human Technopole Early Career Fellowship). The survey respondents therefore reflect a combination of biases, including the professional and social networks of the authors, the inequalities systemic to grant awardees, and those willing to invest significant time to this initiative.

The survey consisted of 106 questions (33 optional) to evaluate the handbook (Figure 3-5), which was made available via Github (see Resources). For each of the 30 commitments, participants were asked to rate its importance from 1 (useless) to 5 (critical), indicate whether it was already established in their research environment, and provide further optional comments. For a subset of commitments (7/30) participants were asked whether the object of the commitment should be *publicly* or *internally* documented. At the conclusion, participants were asked if they would implement, or want their group leader to implement, the Handbook (Supplementary Table 3).

A pilot survey was trialled with the 2024 SAFE Labs workshop participants. This served to revise elements of the handbook and survey that might have caused confusion prior to public release, and identified commitments without a clear consensus for *public* vs *internal* documentation, to be further interrogated through the public survey (7 commitments, Supplementary Figure 1h).

### SAFE Labs Handbook Paid Survey

The paid survey was implemented through Prolific software (https://www.prolific.com/). To maximise similarity to the voluntary data, participants were limited to biosciences PhD students from countries that featured prominently in the volunteer sample. Costs were minimised by selecting 10 of the 30 commitments (with an additional attention-check) that received a range of scores by voluntary PhD students (Supplementary Table 4). Participants were asked to score the importance of these commitments, but did not receive as much background information about the handbook, in order to limit costs. A total of 64 paid respondents were included in analyses (Figure 4-5).

## Data analysis and statistics

Survey data were pre-processed to classify respondents to standardised research roles and categories. For research roles we used the following classification:

| STANDARDISED ROLE | OPEN SURVEY ANSWER |
| --- | --- |
| Group Leader | Assistant Professor, Principal Investigator, Project Leader, Group Leader |
| Postdoc | Researcher, Newly Graduated as PhD, Research Fellow, Research Scientist, Postdoc |
| PhD Student | Intern, Master Student, PhD Student |
| Staff | Research Engineer, Research Assistant, Technician, Lab manager |

For research fields we used the following classification:

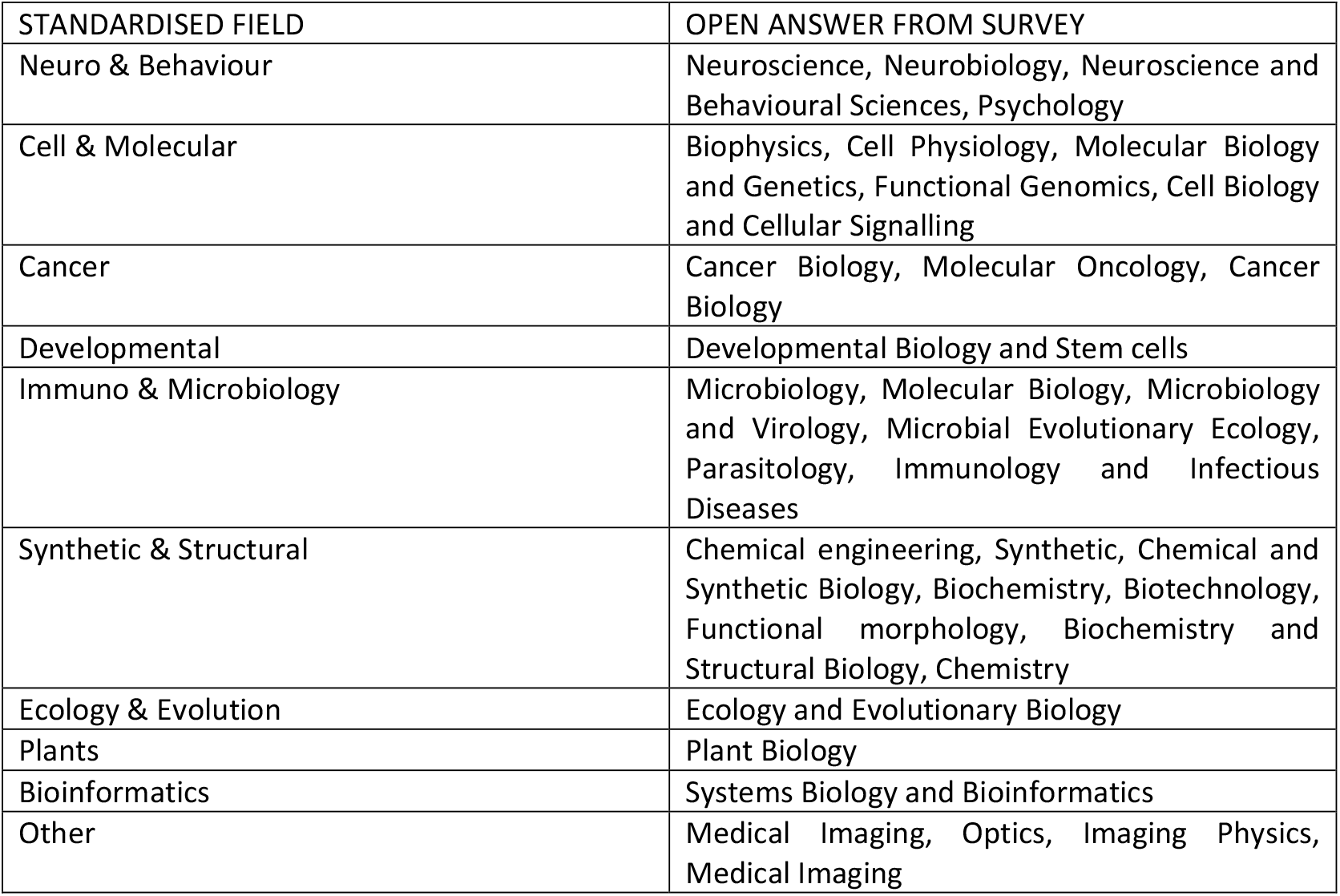

Data were then analysed in MATLAB with custom code. Differences in responses across categories were assessed using the Kruskall-Wallis test or Mann-Whitney U test. The relation between importance and established score was fit with robust linear regression.

## Resources and data availability

The SAFE lab Handbook website: https://safelabs.info/home/safe-labs-handbook/

The SAFE lab handbook repository: https://github.com/SAFE-Labs-Docs/Lab-Handbook

The source code for all analyses: https://github.com/SAFE-Labs-Docs/2025_bioRxiv

## Supplementary materials

Supplementary table 1: SAFE Labs 2024 application form data

Supplementary table 2: SAFE Labs 2024 exit survey data

Supplementary table 3: Handbook public survey data

Supplementary table 4: Prolific paid survey data

## Notes

https://github.com/SAFE-Labs-Docs/Lab-Handbook

https://github.com/SAFE-Labs-Docs/2025_bioRxiv

### Competing Interest Statement

The authors have declared no competing interest.

### Summary of Updates

Minor revisions to the text and figures, including some additional analysis and corrections of minor formatting errors.

