## Supplementart Figures for "The SAFE Labs Handbook: Community-Driven Commitments for Group Leaders to Improve Lab Culture"

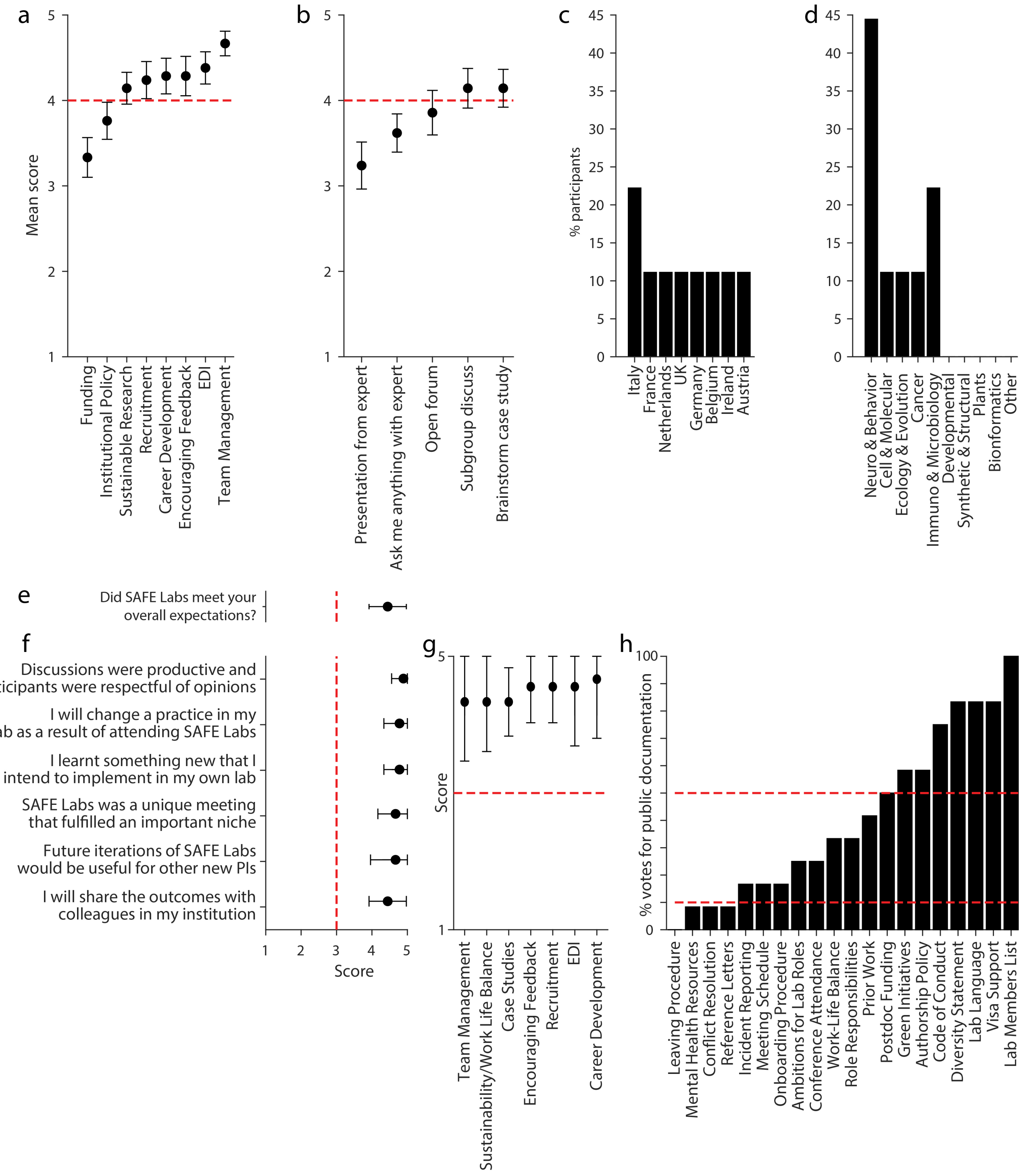


***Supplementary Figure 1. 2024 SAFE Labs Workshop Demographics and Feedback.*** *(**a**) Preference score for proposed SAFE Labs workshop topics, averaged across all applicants, sorted from the least preferred to the favorite. Only topics and formats with high rating (red dotted line) were chosen. Error bars mean ± s.e.m, n = 21 (**b**) As in (a), but for the proposed formats of discussion at the workshop. (**c**) Percentage of selected workshop applicants (n = 9) as a function of country of employment. (**d**) As in (c) for research field. (**e**) Evaluation of the workshop outcome, averaged across participants: range from 1 (= below expectation) to (5 = exceeded expectations). (**f**) Evaluation of workshop feedback statements, averaged across participants: range from 1 (= Strongly Disagree) to 5 (= Strongly Agree). All participants indicated a high rating (i.e. above the red dotted line). (**g**) Rating of the workshop discussion topics, averaged across participants: range from 1 ( = Poor) to 5 (= Excellent). (**h**) Fraction of participants in favor of publicly documenting vs internally documenting each SAFE Lab Handbook commitment. Divisive commitments (within dotted interval) were re-assessed in the public survey (n = 12). For panels e-g, n = 13.*

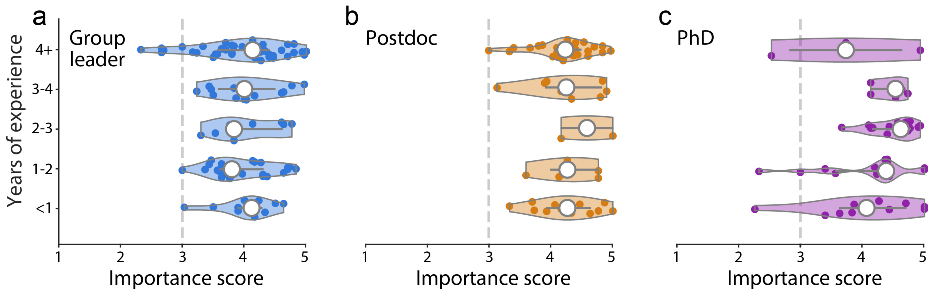


***Supplementary Figure 2. Importance score split by seniority in career stage.*** *(**a**) Distribution of importance rating averaged across commitments for Group Leaders sorted by seniority in their career stage. (**b**) As in (a), but for Postdocs. (**c**) As in (a) but for PhD students.*

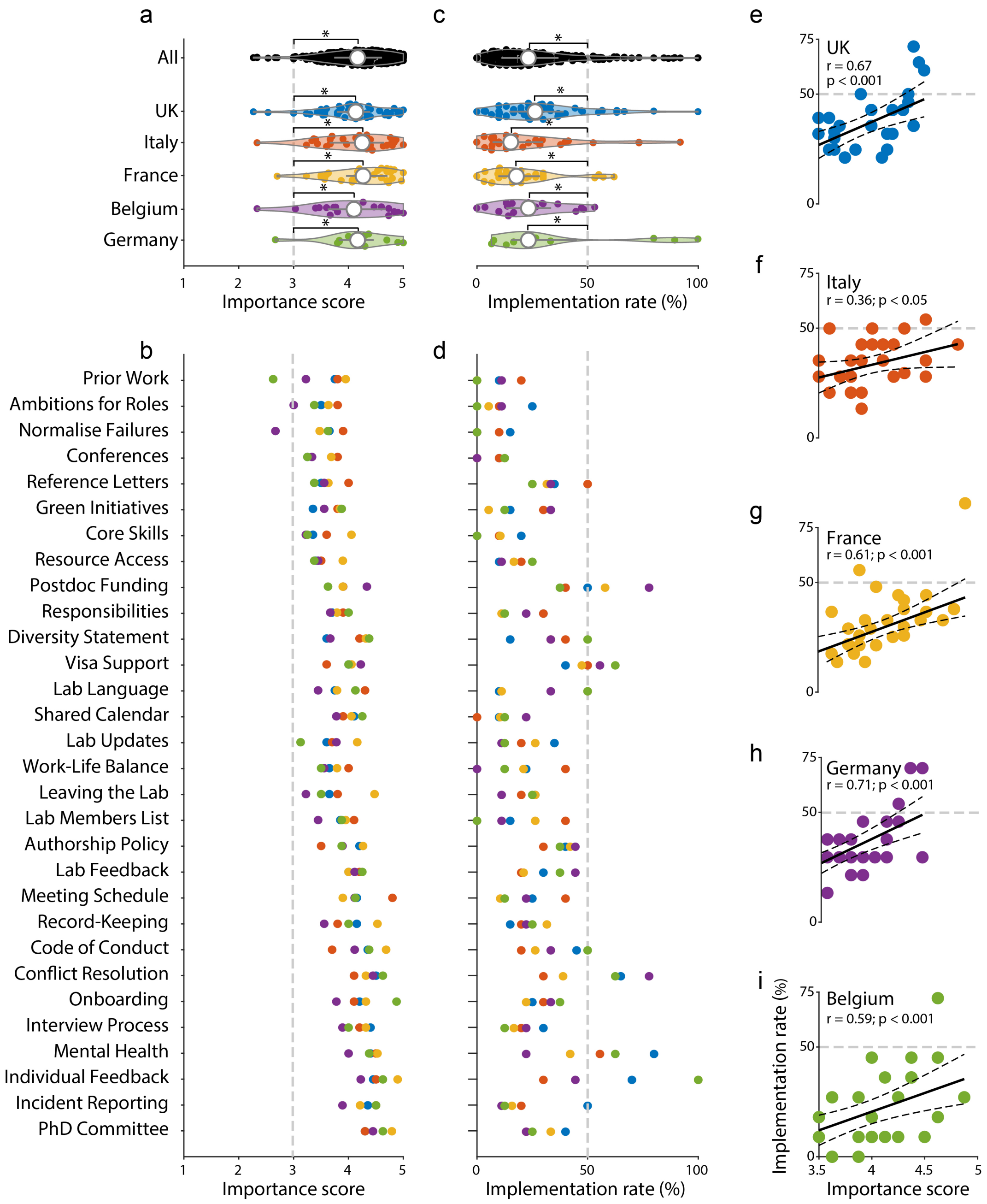


***Supplementary Figure 3. Survey results were comparable across countries.*** *(**a**) Distribution of importance rating averaged across commitments for respondents sorted by the five most numerous employment countries: UK (blue), Italy (orange), France (yellow), Germany (purple), Belgium (green). All countries rated the commitments as significantly important (i.e. above gray dotted line, $*$ = $p < 0.01$) but were not significantly different from each other (Kruskal-Wallis test, $p>0.05$) (**b**) Average importance score for each commitment, sorted as in Figure 4b, for the five main respondent countries. (**c**) As in (a), but for the implementation rate. Respondents from all countries reported similarly low levels of implementation (Kruskal-Wallis test, $p>0.05$), significantly below 50% (gray dotted line , $*$ = $p < 0.01$). (**d**) As in (c), but for implementation rate. (**e**) Relationship between mean importance and mean implementation rate for each commitment as scored by group leaders ($n$ = 20) from the UK. The significant correlation was captured by a robust linear fit (black, dotted confidence intervals, $p<0.001$) (**f**) As in (e), but for Italy (blue, $p<0.05$, $n$ = 10). (**g**) As in (e), but for France (yellow, $p<0.001$, $n$ = 19). (**h**) As in (e), but for Germany (purple, $p<0.001$, $n$ = 9). (**i**) As in (e), but for Belgium (green, $p<0.001$, $n$ = 8).*

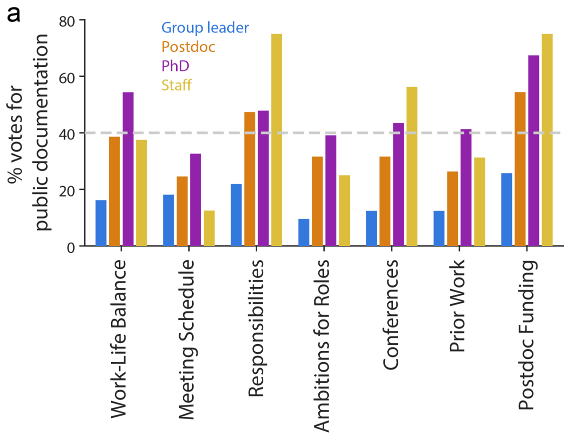


***Supplementary Figure 4. Support for public documentation of selected commitments.*** *(**a**) Fraction of respondents in favor of publicly documenting vs internally documenting selected commitments. n = 105 Group Leaders, 57 Postdocs, 46 PhDs, and 16 Staff.*

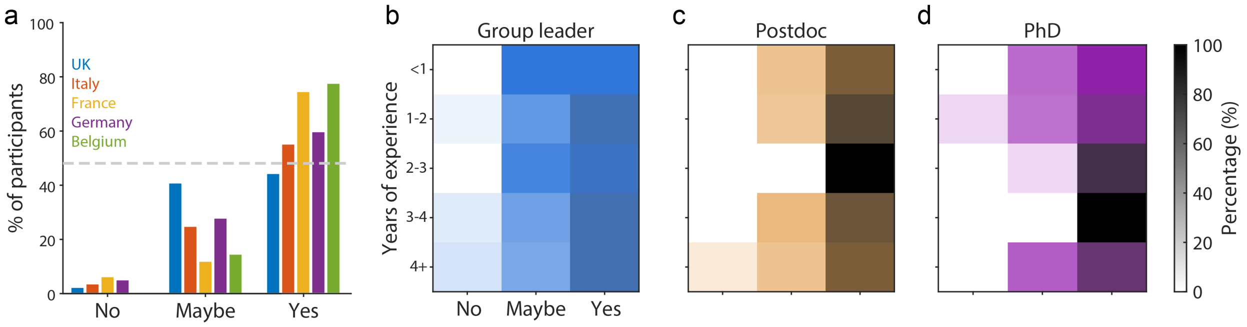


***Supplementary Figure 5. Support for handbook implementation divided by subgroups.*** *(**a**) Fraction of respondents for the main survey in favor of implementing the handbook in their environment split by different countries. (**b**) Fraction of Group Leader respondents (n = 105) for the main survey in favor of implementing the handbook in their environment, split by seniority. (**c**) As in (b) but for Postdocs (n = 57). (**d**) As in (b) but for PhD students (n = 46).*
